# Axial encoding schematics of neural representations of 3D space in freely navigating goldfish

**DOI:** 10.1101/2022.07.07.499255

**Authors:** Lear Cohen, Ehud Vinepinsky, Opher Donchin, Ronen Segev

## Abstract

Navigation is one of the most fundamental cognitive skills for the survival of fish, the largest vertebrate class, and almost all other animal classes. A critical component of the neural basis of navigation is the encoding of space in the activity of single neurons. To study this basic cognitive component in fish, we recorded the activity of single neurons in the central area of the goldfish telencephalon while the fish were freely navigating in a 3D water tank. We found neurons with firing patterns that gradually increased or decreased along spatial axes distributed in all directions. Some of these cells exhibited beta rhythm oscillations. This type of axial coding for spatial representation in the brains of fish is unique among space encoding cells in vertebrates and provides insights into spatial cognition in this lineage.

---

For most animals, the ability to locate themselves in the environment is crucial for survival (1–4). This ability requires the encoding of information about self-position and locomotion (5, 6). Studies in navigating mammals have identified several types of neurons that encode self-position and locomotion in the hippocampal formation (5–9). The most prominent spatial cell type in the hippocampus is the place cell (10), which is activated once the animal is present in the cell’s preferred place field. In addition, other spatial cell types have been identified in the entorhinal cortex and other areas connected to the hippocampus. These include grid cells (11) which are activated in multiple locations spanning the environment, boundary-vector cells which encode proximity to allocentric boundaries (12), and head direction cells (13).

In the goldfish lateral pallium, a possible homologue of the mammalian hippocampal formation (14), a recent study described the existence of single cells that encode the edges of the environment, the fish’s head direction, speed, and velocity (15). However, there was no evidence of neuronal activity in a specific place field, although this has been extensively documented in mammals (16) and recently also reported in birds (17, 18). In addition, there was no evidence of rhythmic neural oscillations associated with place encoding, unlike what has been found in the theta frequency range of the mammalian hippocampal formation (19).

To better understand the fundamentals of spatial cognition, a comparative approach can shed light on the universality of the mammalian model in all vertebrates or whether different classes have developed different computational schemes. For this purpose, it is crucial to determine how information on position is represented in the brain of fish, the largest vertebrate class. Since water is significantly denser than air, fish are subjected to a steeper pressure gradient along the vertical axis. This might imply that information about position in 3D space is encoded in the fish brain differently than the locally activated cells that were shown to exist in 3D environments in bats (20) and rats (21).

To probe navigation in fish, we recorded multiple single-cell activity from the central area of the goldfish telencephalon. This brain region is thought to integrate information from multiple neighboring pallial regions (22–24). We hypothesized that it would integrate representations of space in the fish telencephalon. During the recordings, the goldfish could freely explore an experimental water tank including along its vertical and horizontal axes.

## Results

To understand how 3D space is represented in the teleost brain, we measured single-cell activity in the central area of the telencephalon of freely behaving goldfish while they explored a 3D water tank (Figure 1). We first trained the fish to swim continuously in a rectangular water tank (measuring 0.7 m in height, 0.2 m in length and 0.7 m in width, Figure 1A) by feeding them in various places in the tank. After the fish became familiar with the water tank and had adapted to exploring its entire environment freely, we implanted an extracellular wireless recording system into its brain (see *Materials and Methods*). We then let the goldfish swim in the water tank while recording neuronal activity and tracking its position. Neuronal activity was recorded extracellularly using tetrodes, and single cells were identified using spike sorting (Figure 1 C-E). Subsequently, we analyzed the associations between cell activity and the fish’s trajectory (see *Materials and Methods*).

**Figure 1.**
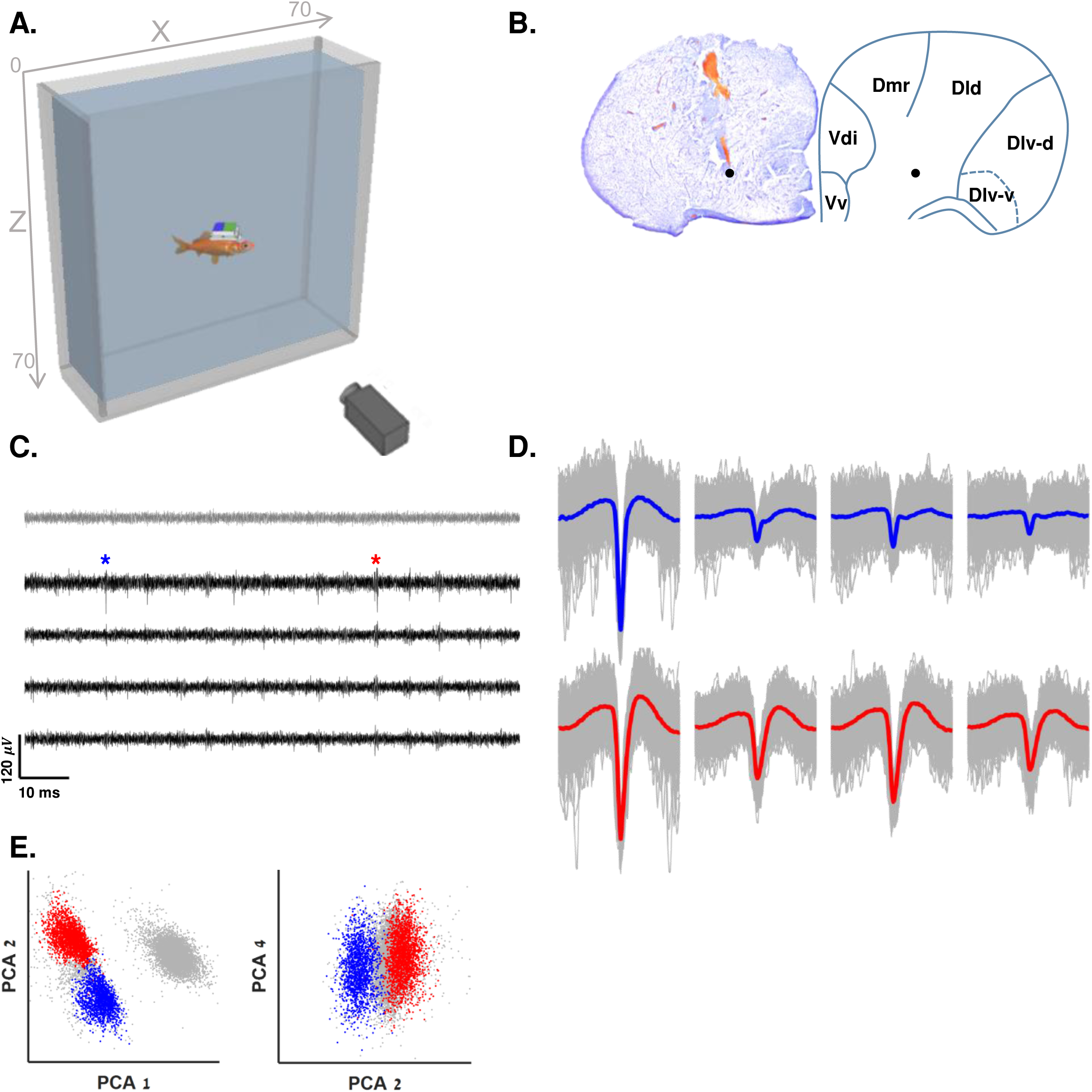
Experimental setup and spike sorting. **A.** Schematic overview of the experimental setup: a fish swims freely in a water tank with electrodes, and a wireless data logger mounted on its head. The fish’s movements are recorded by a Raspberry Pi camera positioned in front of the tank. **B.** Examples of recording site in the goldfish central telencephalon and the corresponding brain region (right panel, anatomical diagram based on (41)). Black dots represent locations in which spatially modulated cells were recorded. **C.** Example of a raw recording from a tetrode (black traces) and a reference electrode (gray) in the fish’s central telencephalon. Neural activity can be seen in the tetrode alone. Blue and red asterisks correspond to the blue and red clusters in panels D-E. Blue cluster forms the cell presented in Supplementary Figure 1C. Red cluster forms the cell in Figure 2 A-D. **D.** Waveforms of the two neurons after spike sorting. **E**. Projections on the main principal components of the data from the tetrode of all spike candidates that crossed the threshold. Other clusters were not distinguishable from other multiunit activity and neural noise.

The results showed that for a considerable portion of cells (∼30%), neural activity was modulated by the fish’s position in the environment (see examples in Figure 2). In some neurons, activity was modulated along a preferred principal axis in the environment. For example, Figure 2 A-D and Supplementary Video 1 show a cell with a firing rate that was modulated along the horizontal axis. This cell’s neural activity gradually decreased with the distance of the fish from the left wall of the water tank. This was manifested in the spiking activity (red dots, Figure 2A) throughout the fish’s trajectory (black curve) as well as in the occupancy-corrected rate map in the water tank (Figure 2B).

**Figure 2.**
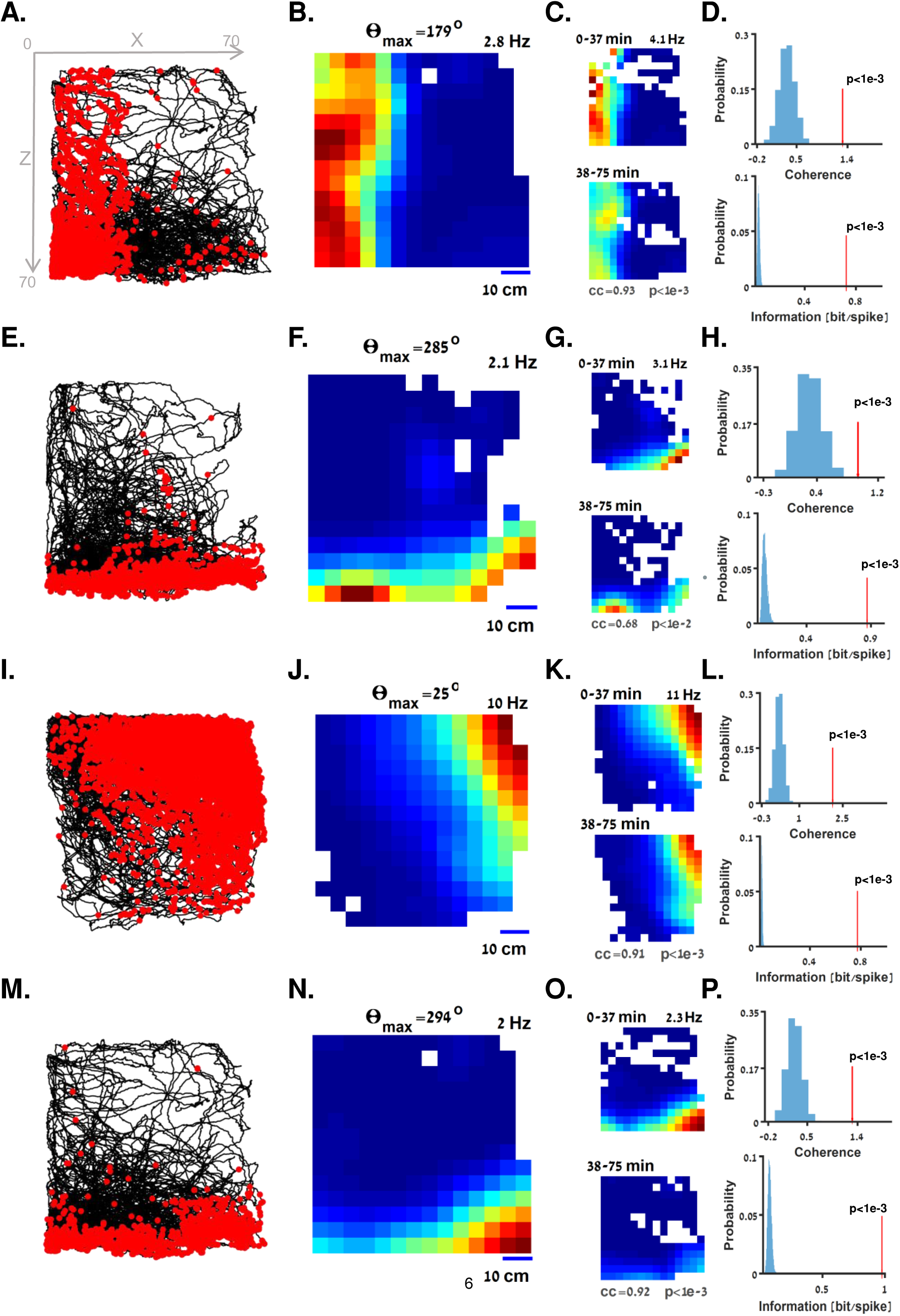
Spatially modulated cells in the goldfish brain. **A-D.** Example of a spatially modulated cell with preferred tuning to the horizontal axis (experimental setup is shown in Figure 1A). **A.** The fish trajectory (black curve) is presented together with the location of the fish when each spike of a single cell occurred (red dots). The neuron was mainly active when the fish was near the left wall of the tank. **B.** Firing rate heatmap of the cell in A, color coded from dark blue (zero firing rate) to dark red (maximal firing rate, indicated on the upper right side of the panel). **C.** In-session stability of the cell in A. The similar rate of the cell in the first (top panel) and second (bottom panel) halves of the experiment suggest stable activity within the recording session (correlation coefficient and p value are indicated, see *Materials and Methods*). **D.** Spatial coherence (red arrow, top panel) and information (red arrow, bottom panel) values of the cell in A are higher than those of 5000 shuffled spike trains obtained from the same dataset (blue histograms, see *Materials and Methods*). **E-H.** Example of a spatially modulated cell with preferred tuning to the vertical axis. **I-P.** Additional examples of spatially modulated cells with gradual tuning to position along other axes of space.

To test statistically whether this neuron encoded components of position, we tested for in-session stability and for spatial coherence and information. The similarity between the firing rate maps for the first (0-37 min) and the second (38-75 min) halves of the session indicated in-session stability (Figure 2C, correlation coefficient=0.93, p<1e-3, see *Materials and Methods*). The cell’s spatial coherence (Figure 2D, top panel, red arrow) and information (Figure 2D, bottom panel, red arrow) were greater than the corresponding values for 5000 shuffled spike trains (blue histograms, see *Materials and Methods*).

Another cell tuned to position along the vertical axis is presented in Figure 2 E-H. Additional examples of cells with graded firing patterns along one of the major axes of space (the horizontal and vertical axes) are presented in Supplementary Figure 1 and Supplementary Figure 2 A-D, as well as in Supplementary Videos 2 and 3.

Not all cells were tuned to the major axes of space, but rather to a specified minor axis. An example of this type of cell is presented is Figure 2 I-L. For this cell, firing was correlated to the distance from the top right corner of the water tank. Other examples of cells modulated along minor axes of space are presented in Figure 2 M-P and in Supplementary Figure 2 E-G.

The spatial tuning properties of the cells shown in Figure 2 are presented in Figure 3. To further assess the tuning of the cells to the spatial axes, we calculated the correlation coefficient of the cell activity and the fish’s position on each axis (Figure 3A, red dot, which corresponds to the cell presented in Figure 2 A-D) and compared it to the 2D correlation coefficients of 5,000 shuffled spike trains (grey dots, the 95^th^ percentile is depicted) generated using an ISI shuffling procedure (see *Materials and Methods*). This cell was correlated to a position in the negative X-axis direction (*Θ*_*max*_ = 179^*o*^, see *Materials and Methods*). This was further confirmed by the spiking activity of the cell (red dots, Figure 3B) over the projection of position along the preferred axis (black curve). This cell’s tuning curve of firing rate to position along the *Θ*_*max*_ axis showed a clear pattern (Figure 3C, blue curve), whereas no clear tuning pattern was visible for the orthogonal axis (*Θ*_*max*_ + 90^*o*^, orange curve). Similar results are presented in panels D-F, G-I and J-L and correspond to the cells presented in Figure 2 E-H, I-L and M-P, respectively.

**Figure 3.**
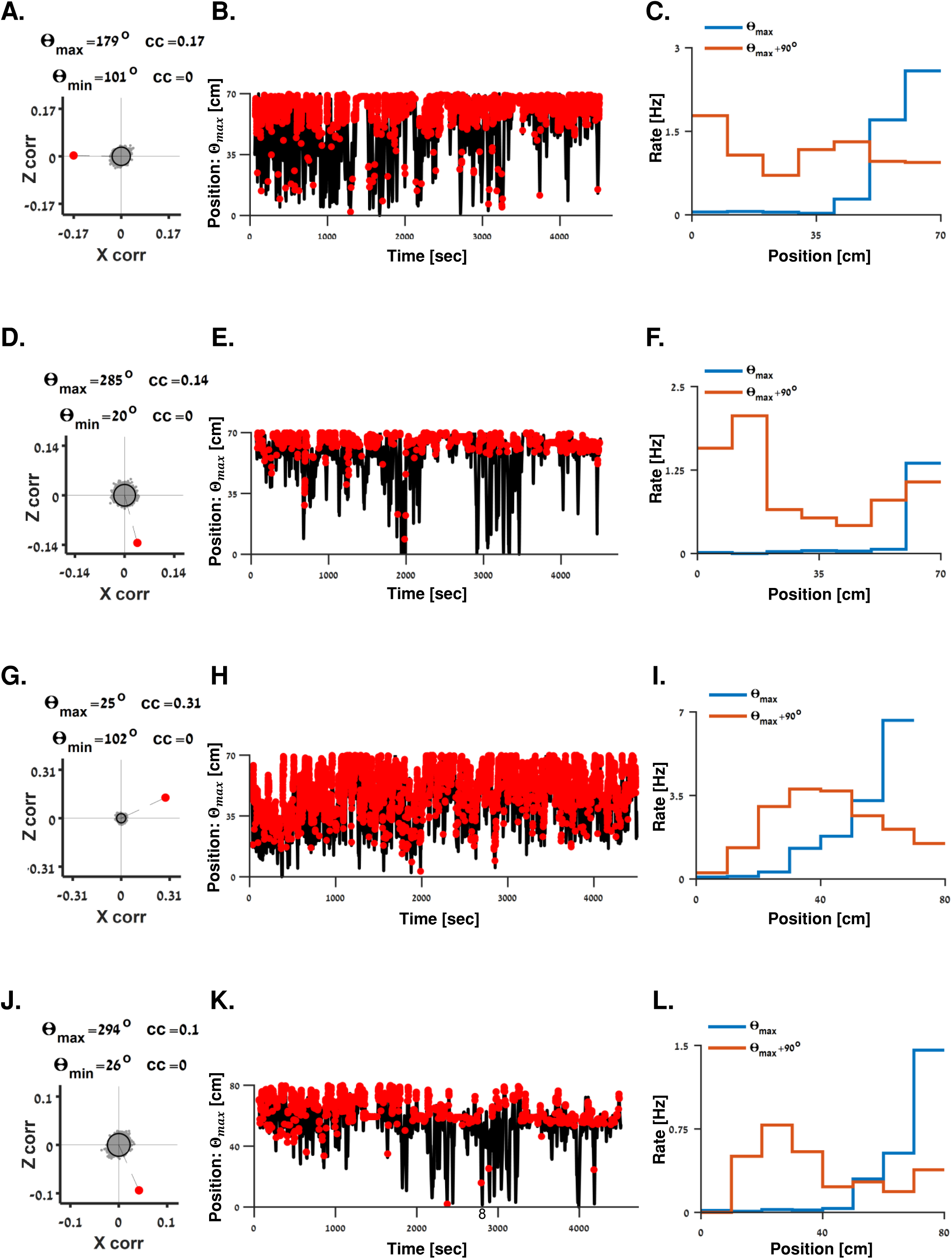
Tuning properties of spatially modulated cells. **A-C.** Tuning properties of the cell presented in Figure 2 A-D. **A.** 2D correlation coefficient of firing rate and the fish’s position of the cell (red dot) and 5000 shuffled spike trains (gray dots, the 97.5 percentile is depicted), suggesting this cell is a correlated to a position in the negative X direction. Preferred and null tuning directions (see *Materials and Methods*) and correlation coefficients (*cc*) are indicated. **B.** Spiking activity (red dots) superimposed with the position of the fish (black curve) along the preferred axis. **C.** Tuning curve of the cell’s firing rate to the position of the fish along the preferred axis (*Θ*_*max*_, blue curve) and its orthogonal axis (*Θ*_*max*_ + 90^*o*^, orange curve). Gradual tuning is shown for the *Θ*_*max*_ axis. **D-L.** Tuning properties of the cells presented in Figure 2 E-H, I-L and M-P, respectively.

To differentiate the axial effect from a simple border effect, we conducted a control experiment in which we recorded the neural activity of spatially modulated cells before and after a geometric change induced in the environment. Examples of these cells are presented in Figure 4. In one example, the cell’s neural activity gradually increased with the fish’s position along the positive vertical direction (Figure 4 A-C). After adding a step covering half of the bottom of the water tank (Figure 4D), the cell’s activity pattern (Figure 4 E-F) remained mainly near the bottom of the water tank. Despite the major difference in the geometry of the environment, similar tuning curves of rate to position along the preferred axis of this cell in the first (Figure 4G, blue curve) and second (orange curve) recording sessions were observed (correlation coefficient=0.87, p=0.01, see *Materials and Methods*). We verified the cell’s identity in the electrophysiological recordings by tracking the cell’s mean spike waveforms across the two sessions (Figure 4H).

**Figure 4.**
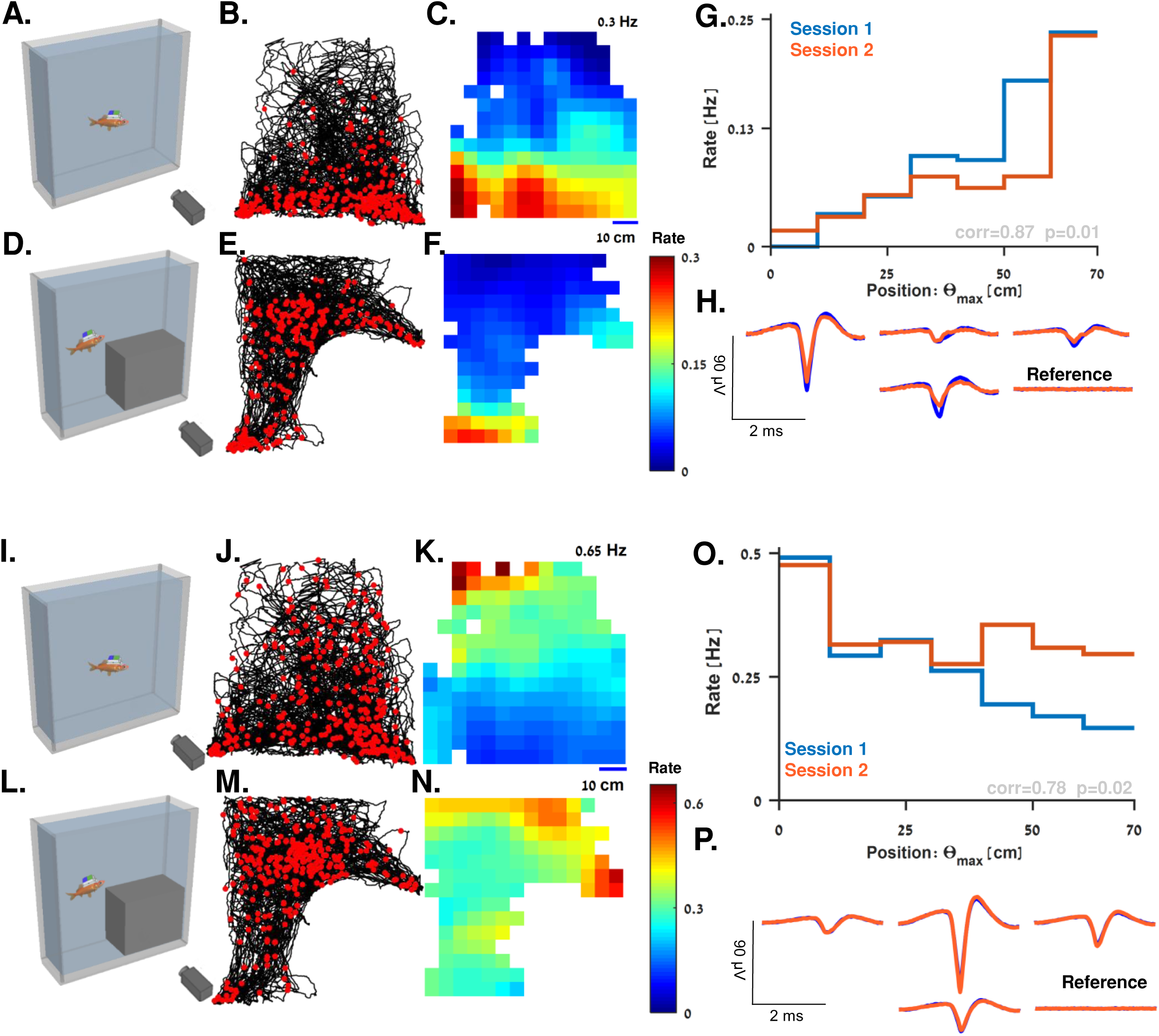
Changing environmental geometry. **A.** The first session was recorded in the main experimental water tank. **B-C.** Example of a spatially modulated cell with tuning to the positive vertical axis. **B.** The fish trajectory (black curve) is presented together with the location of the fish when each spike of a single cell occurred (red dots). The neuron was mainly active when the fish was near the bottom of the tank. **C.** Firing rate heatmap of the cell in B, color coded from dark blue (zero firing rate) to dark red (maximal firing rate, indicated on the upper right side of the panel). **D.** A step was inserted into the water tank to modulate the geometry of the environment. **E-F**. Fish trajectory and the heatmap of firing rate over the environment indicates that the firing rate increased with position along the vertical axis. **G.** Tuning curve of the cell’s firing rate to position of the fish along the vertical axis in the first (blue curve) and second (orange curve) recording sessions. In both sessions, the rate gradually increased with position along *Θ*_*max*_. **H.** Mean spike waveforms in the different sessions (blue-first session, orange-second session) indicate this is the same unit. **I-P**. Another example of a spatially modulated cell whose firing rate decreased with position along the vertical axis in both sessions.

Another cell tuned to position in the negative vertical direction is presented in Figure 4 I-P. This cell also showed stable tuning (correlation coefficient=0.78, p=0.02, see *Materials and Methods*) across two sessions with a different geometry. The similarity of the tuning curves between sessions suggests these cells are not activated by a simple border tuning mechanism.

Population analysis showed that 31 out of the 101 recorded cells, i.e., about 31% of all the well-isolated units recorded in the 14 fish used in this study exhibited spatial modulations, i.e., passed the statistical tests for in-session stability, spatial coherence and information. Of the 31 spatially modulated cells, 4 cells had a diffuse firing pattern and were removed from the axial tuning analyses.

### Axial coding of space

To demonstrate axial encoding of space, we needed to confirm that the spatially modulated cells could be better described by axial tuning than by simple place encoding. For this purpose, we tested two tuning models: 1. Sigmoid tuning along the cell’s preferred axis drawn from the spatial modulation pattern and 2. Gaussian tuning in the form of a two-dimensional Gaussian centered proximal to the environmental boundaries.

We fitted the two models for the firing rate map of each cell and tested the goodness of fit for the two models. This process was first validated using a simulated dataset (examples are presented in Supplementary Figure 3, see *Materials and Methods*). Using the simulated data, we set the thresholds (dashed lines in Figures 5 A-B, see *Materials and Methods*) for classification of a cell as axial encoding or Gaussian encoding (as in the form of a place cell).

**Figure 5.**
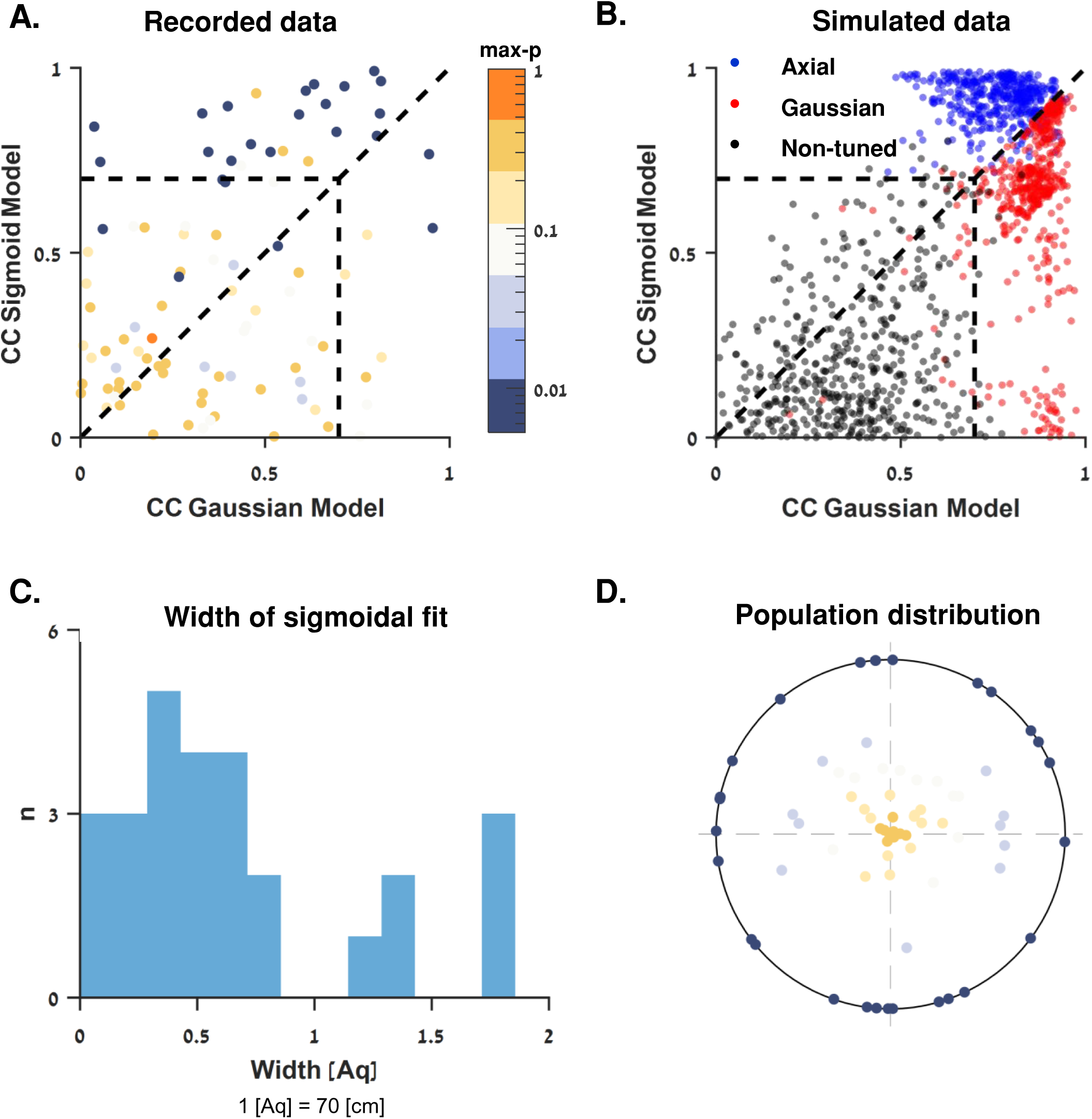
Population analysis of spatially modulated cells. **A.** Correlation coefficients between recorded data rate maps and the rate maps generated by a Gaussian model and a sigmoid model (see *Materials and Methods*) suggesting that the spatially modulated cells encode position using axial encoding schematics. Thresholds (dashed lines) were derived from a simulated dataset (panel B). The color bar on the right-hand side of panel A spans the population max-p values (see *Materials and Methods*) on a logarithmic scale and shows the distribution of cells in panels A and D. **B.** To validate the classification in A, three groups of 500 firing patterns were simulated (corresponding to the examples presented in Supplementary Figure 3), and correlation coefficients were calculated for each fitting method (see *Materials and Methods*). The three groups-axial encoding, Gaussian encoding, and non-encoding could clearly be classified using boundaries of 0.7 and above or below the identity line. **C.** Distribution of widths for the sigmoid curves fitted to the spatially modulated cells relative to the water tank’s width. Units: 1 [Aq] = 70 [cm]. **D.** The population max-p values and preferred direction *Θ*_*max*_ (see *Materials and Methods*) of all 101 single units recorded from the brains of 14 fish. The dots are distributed on a radial plot where 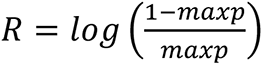 folded to the range [0, 5] is the radius and *Θ*_*max*_ is the angle.

The results showed almost all the spatially modulated cells had a better fit to the sigmoid model than to the Gaussian model (see Supplementary Table 1). This emerged in the correlations between the data and the models (Figure 5A, dark blue dots represent the cells with the strongest spatial modulations). Specifically, the activity of the spatially modulated cells gradually increased or decreased with the position of the fish along the cell’s preferred axis, regardless of its position along the orthogonal axis. This lends weight to the claim that spatially modulated cells have a preferred axis of tuning.

In addition, the widths of the sigmoid curves fitted to the spatially modulated cells were distributed around 67% ± 53% (Mean ±SD) of the water tank’s width (Figure 5C). This fit differs from a simple border cell which is expected to have sharp sigmoidal tuning near a specific boundary in the environment (25).

We did not find clear groups of preferred directions for the spatially modulated cells. Figure 5D shows the distribution of Θ_*max*_ and the max-p values (see *Materials and Methods*) of the entire recorded population. While the cells tuned along the vertical axis appear to be more abundant, there was not enough data to investigate this further and future study is needed to clarify this point.

### Beta oscillations in spatially modulated cells

Some of the spatially modulated cells exhibited rhythmic neural activity. An example of one such cell is presented in Figure 6 A-D. This cell is spatially modulated by axial tuning along the vertical axis (Figure 6 A-B). Examining the histogram of inter-spike interval of this cell (Figure 6C) revealed periodic spacing between the peaks of the histogram. This was also shown by calculating and plotting the power spectral density function of the histogram in C (Figure 6D), where a local maximum appeared at around 16Hz: in other words, this neuron oscillated rhythmically in the low-beta frequency range. A counter-example of a cell that was not tuned to space is presented in Figure 6 E-H.

**Figure 6.**
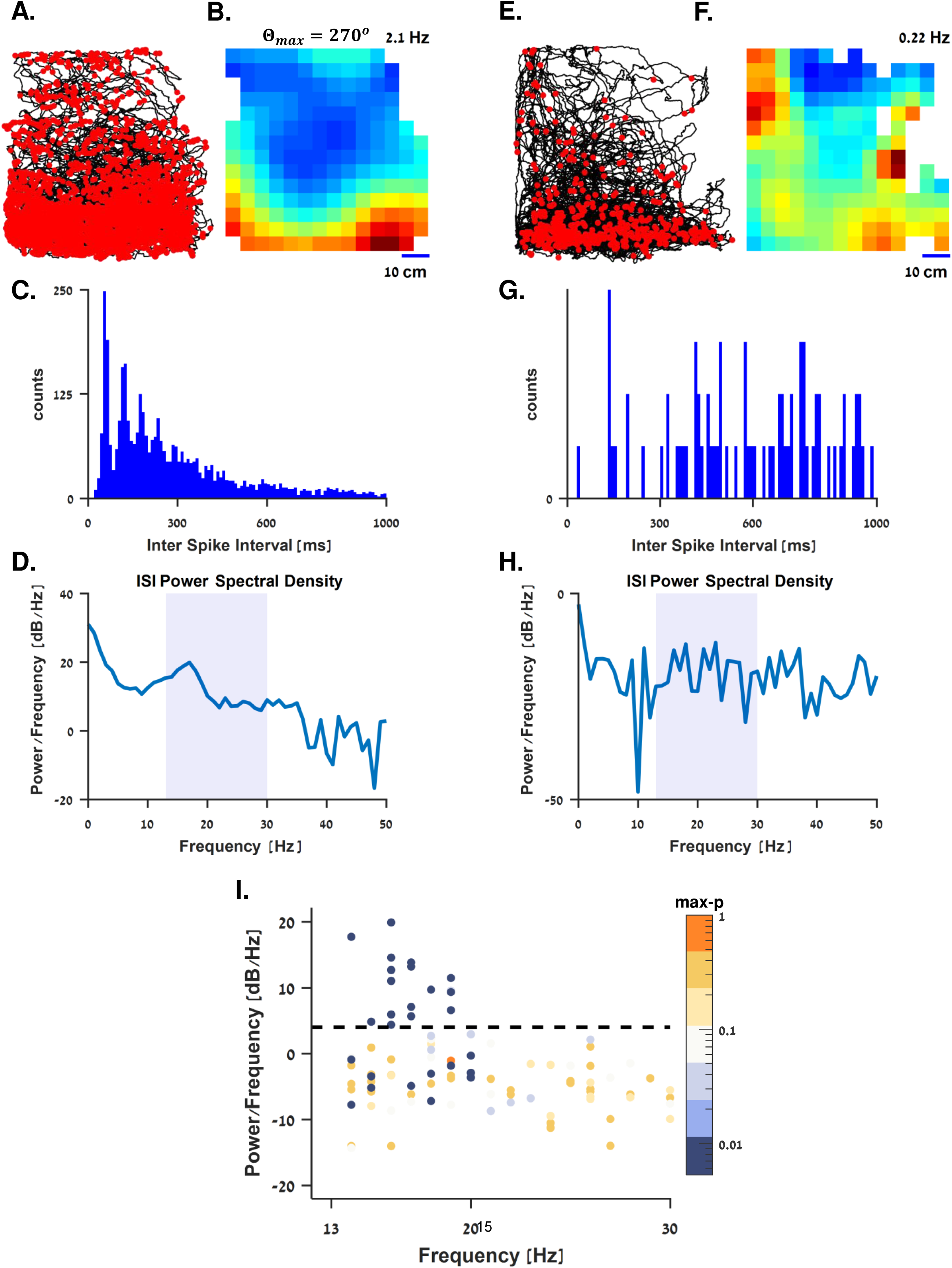
Beta oscillations in spatially modulated cells. **A-D.** Example of a spatially modulated cell with periodic oscillations of the inter-spike-interval (ISI). **A.** The fish trajectory (black curve) and spike locations (red dots) of a cell tuned to the vertical axis. **B.** Firing rate heatmap of the cell in A, color coded from dark blue (zero firing rate) to dark red (maximal firing rate, indicated on the upper right side of the panel). **C.** Inter spike interval histogram of the cell in A. Even spacing between the peaks suggests periodic oscillations of the neural activity. **D.** Power spectral density of the histogram in C. Colored background marks the typical range of beta waves (13-30 Hz). A local peak is shown at 16 Hz suggesting this cell exhibited beta-rhythm oscillations. **E-H**. A counter-example of a cell with no clear space tuning and no specific pattern in the ISI histogram. **I.** Maximal power spectral density in the beta waves range for the entire population. The color bar spans the population’s max-p values (see *Materials and Methods*) on a logarithmic scale. As shown, roughly half of the spatially modulated cells (dark blue dots) exhibited beta rhythm oscillations around 17 Hz.

Examining the position and magnitude of the peak power spectral density in the beta range of the entire population (Figure 6I, color-coded by the max-p index, see *Materials and Methods*) revealed a clear connection between the beta oscillations and the spatially modulated cells in the central telencephalon of the goldfish. To estimate the properties and prevalence of the beta oscillations in the population, we used a threshold of 4 dB/Hz for the peak power of the spectral density. The findings showed that 16 of the 27 spatially modulated cells (∼59%) and 1 out of the other 70 cells crossed this threshold with a peak spectral density in 17±1.53 Hz (mean±standard deviation).

### Stability of the analysis

To ensure that our space tuning analysis was statistically valid even when the swimming trajectory did not cover the water tank homogeneously (Supplementary Figure 4), we ran our analysis pipeline on a set of simulated data. The simulated cells’ activity was drawn from three different tuning curve types (500 shuffled spike trains for each tuning curve) over a fish’s recorded trajectory which did not cover the water tank homogenously (corresponding to the trajectory of the cell presented in Supplementary Figure 1B). We used tuning curves from three different fish that gradually decreased (Supplementary Figure 4A, right panel), were quasi-uniformly distributed (Supplementary Figure 4B, right panel), or gradually increased (Supplementary Figure 4C, right panel) with the position of the fish along the vertical axis (see *Materials and Methods*).

For the decreasing axial tuning, 499 of the 500 simulated cells were indeed classified as spatially modulated. This was also true for 498 out of the 500 cells obtained for increasing axial tuning (false positive rate<0.004, Supplementary Figure 4D). None of the cells obtained from the quasi-uniform firing rate (Supplementary Figure 4D) were classified as spatially modulated (false negative rate<0.002). This confirms that the analysis for space tuning was not sensitive to occupancy correction artifacts in the firing rate map due to partially visited parts of the water tank. Finally, to verify that these cells did not encode other aspects of locomotion such as swimming speed, we controlled for the swimming speed distribution across the water tank to make sure no clear gradual pattern emerged (Supplementary Figure 5).

## Discussion

The analyses provide evidence that the activity of neurons in the central area of the goldfish telencephalon is monotonously correlated to a specific spatial axis. The axial code could be tuned to all directions of space. This encoding scheme differs from the local encoding of mammalian place and grid cells, which was also shown to be present in 3D environments in rats (21) and bats (26). It also differs from the typical mammalian border cell in which the firing rate decreases sharply near one or more of the environmental boundaries (25). This encoding scheme of space may thus constitute a new building block in the navigation system of vertebrates generally, or aquatic animals specifically.

These findings contribute to efforts to define the basic inventory of spatial and kinematical cells in the goldfish telencephalon and strengthens claims that the telencephalon plays a crucial role in representations of space and locomotion during navigation tasks. The abundance of spatially modulated cells in the central part of the telencephalon underscores the importance of this brain region for navigation. For these reasons, the data reported here constitute an important step towards a better understanding of the representation of space in the brains of fish, the largest group of vertebrates. Using a comparative approach to the well-studied mammalian navigation system can thus help decipher which parts of this system have evolved in different taxa and how.

It is possible that a subset of the cells presented here reflected depth encoding. Several studies have suggested that fish can sense hydrostatic pressure in the telencephalon and use it for navigation (27–37). However, additional experiments are needed to show depth encoding *per se*. For example, neural activity could be recorded in an environment that changes in hydrostatic pressure in an invariant visual scene.

Our findings are different from the previously reported edge cells observed in the goldfish Dlv region in a shallow water tank (15). Edge cells in that study were activated when the goldfish swam near any of the environmental borders. Here, by contrast, most of the spiking activity gradually decreased with distance from a specific boundary or even part of a boundary. One possible explanation is that in the current study the fish also navigated in the depth dimension, which made the sensory cues in the experimental setup more like the cues it uses to navigate in its natural habitat. Another explanation is related to differences in the sites of the sub regions of which neural activity recorded from in the two studies.

Alternatively, the neurons we recorded may have encoded the distance from the environmental boundaries or from salient cues in the environment. In other words, they could have encoded the bottom of the water tank, the water’s surface, or the walls or corners of the water tank. This type of encoding has been argued to be integral to place encoding in the rat and was ascribed to what were termed “boundary vector cells” (12). Their activity was shown to gradually decrease with distance from an allocentric boundary, regardless of geometric changes induced in the environment. To further test this hypothesis, additional experiments including geometric changes in the environment are needed.

Our results suggest the presence of a population of spatially modulated cells with gradually modulated firing patterns along different axes of space. Thus, the mechanism underlying this phenomenon may be the gradual encoding of space. Recording single neurons in the goldfish central telencephalon while it is swimming in a larger environment could help further test this hypothesis.

In addition, we observed neural oscillations in the low-beta range (13-20 Hz) in the activity of many of the spatially modulated cells and almost none of the cells that were not modulated by position. Neural oscillations associated with spatial memory are often reported to be in the theta frequency range (19). We did not find evidence in the literature for an association between spatial localization and beta oscillations, and further investigations are needed to establish this point.

Thus overall, this study contributes to the growing body of knowledge on the representation of space in the telencephalon of fish. Since this is an important group of vertebrates, it is crucial to continue this line of research to better understand how these building blocks lead to successful navigation in fish.

## Materials and Methods

### Experimental model

A total of 14 goldfish (*Carassius auratus*) 13–18 cm in length and weighing 100–250 g was used in this study. Each fish was housed in a home water tank at room temperature and brought to the experimental water tank for the recording sessions. The room was set to a 12/12 h day-night cycle under artificial light. All the experiments were approved by the Ben-Gurion University of the Negev Institutional Animal Care and Use Committee and were in accordance with government regulations of the State of Israel.

### Wireless electrophysiology

The implant preparation and transplanting process, as well as the behavioral electrophysiology details are fully described in our previous publications (38, 39). Briefly, extracellular recordings obtained from the goldfish central telencephalon using 3 or 4 tetrodes were logged on a small recording device (Mouselog-16, Deuteron Technologies Ltd., Jerusalem, Israel) placed in a waterproof case mounted on the fish’s skull. In addition to the tetrodes, a reference electrode was placed near the brain to detect possible motion artifacts rather than neural activity. The tetrodes were moved in the brain between recording sessions using the built-in microdrive of the implant. Wireless communication via a PC and a transceiver (Deuteron Technologies Ltd., Israel) was used to control and synchronize the data logger. The neural signals were recorded at 31,250 Hz and high-pass filtered at 300 Hz. A colored styrofoam marker attached to the waterproof case was used to neutralize the buoyancy of the implant (i.e., total average density of 1 g/cm^3^) and to detect the fish’s position and swimming direction on the video recordings.

### Surgery and stereotaxic procedure

During surgery, the anesthetized fish was placed in a holder on the operating table and perfused through its mouth with water and anesthetic (MS-222 200 mg/l, NaHCO3 400 mg/l 1, Cat A-5040, and Cat S-5761, Sigma-Aldrich, USA). All surgery specifics are described in detail in (39). We targeted the central area of the telencephalon by taking the anterior mid margin of the posterior commissure as the zero point for the stereotaxic procedure (40). Then, using a mechanical manipulator, we moved the tetrodes 1 mm posteriorly and 1.5 mm ventrally. We used the microdrive attached to the tetrodes to further adjust them in the dorsal/ventral axis after surgery.

### Experimental water tank

The experimental water tank was 0.2 m in length X 0.7 m in width X 0.7 m in depth, creating a quasi-3D environment with an extended depth axis and a foreshortened length dimension (Figure 1A). In some of the experiments we artificially altered the geometry of the experimental environment by placing a Perspex step of 0.2 m in length X 0.35 m in width X 0.35 m in height in the experimental water tank to change the geometry of the space where the fish could swim.

### Video recording

A Raspberry Pi camera (Raspberry Pi 3B microprocessor) oriented towards the center of the water tank was used to localize the fish in the width and depth dimensions. To synchronize the neural activity and the video recordings, we used Arduino Uno, which sends simultaneous TTL pulses to the data logger’s transceiver and the Raspberry Pi unit. Each recording session lasted 55-75 minutes while the fish navigated freely in the water tank.

### Histology

The brains of 12 out of 14 fish were fixed in 4% paraformaldehyde overnight (Electron Microscopy Sciences, CAS #30525-89-4), and then were immersed in a 40% glucose solution for cryoprotection. After freezing, the brains were cryo-sliced (40 um slices) and Nissl stained to reveal the position of the electrodes in the brain (i.e., the central area of the telencephalon). All slices were then scanned by automated microscope (Panoramic® MIDI II, 3DHISTECH, Hungary) with a 20x (NA 0.8) and a 40x (NA 0.95) objective. The slices were compared to the neuroanatomical landmarks of (41) to confirm the recording region.

### Spike sorting

The raw neural data were band-pass filtered at 300–7,000 Hz. Then, a threshold was set to detect the action potential timings. Manual single cell clustering was done for each tetrode separately by standard spike sorting methods (42, 43), including PCA analysis of the spike amplitudes, widths, and waveforms across all four electrodes. Units not clearly separated in the PCA space and units unstable over time were eliminated from further analysis. Additionally, spikes that appeared simultaneously in more than one tetrode or reference electrode were considered motion artifacts or electronic noises and were removed from the analysis. Examples of clean spikes are presented in Figure 1 C-E. More details can be found in (38, 39).

### Fish trajectory and firing rate map analysis

The fish’s location and orientation in each video frame were detected using the open-source pose-estimation algorithm by DeepLabCut (44). This deep-learning algorithm detected the coordinates of the colored styrofoam marker attached to the implant on the fish’s head in each video frame. Next, the coordinates were translated to the width-depth position in units of cm. Then, the fish’s trajectory and spike positions were binned using a 5 cm x 5 cm grid to obtain auxiliary maps of occupancy per bin and spike count per bin, respectively. A bin-by-bin division of spike counts by occupancy yielded the occupancy-corrected firing rate map for each cell. The corrected rates in each map were color-coded from zero (dark blue) to the maximal firing rate of each cell (dark red). Bins in which the fish spent fewer than 2 seconds were discarded from the analysis and colored white in the figures.

### Spatially tuning statistics

To evaluate spatial modulation, we applied the standard statistical tests for spatial modulation in the literature: in-session stability, spatial coherence and information (45). To test for in-session stability, we calculated the firing rate maps for the first and second halves of each experiment and calculated the correlation coefficient between them. Then, 5,000 shuffled spike trains were obtained by calculating the inter-spike intervals (ISI), shuffling the inter-spike intervals by random permutation, and using the cumulative sum to obtain a shuffled spike train. For each shuffled spike train, we created a heatmap using the trajectory of the second half of the experiment and calculated the correlation coefficients between the shuffled heat map and the heat map of the first half of the experiment. Comparing the shuffled results with the data yielded a p-value for stability representing the probability of obtaining greater in-session stability than that of the cell by chance.

For the shuffled spike trains, we also calculated the spatial information and coherence values and compared them to the cellular result to obtain two additional p-values. We assigned each cell a max-p value (as used in Figure 5 A and D and in Figure 6I) which was the largest p-value of those calculated in the three tests. Cells with a max-p<0.025 and for which the spatial information was at least 0.1 bit / spike were counted as spatially modulated cells. This category was used solely for the purposes of determining tuning width in spatially modulated neurons and to help estimate the prevalence of beta rhythm and spatial tuning in the population. In the latter case, we also tested different max-p and spatial information thresholds and observed that the prevalence of spatial tuning was not strongly affected (see Supplementary Table 2).

### Spatial correlation and axial preferences

As one measure of neuronal tuning, we calculated the correlation coefficient between the cellular firing rate and the position in both the width and depth axes. We used a shuffle analysis to get a perspective on the magnitude of these correlations. Specifically, 5,000 shuffled spike trains were obtained by ISI shuffling as described above. For each shuffled spike train, the correlation coefficients were calculated as with the original data. The results of this analysis were used solely for graphical assessment (Figure 3 A, D, G and J) whereas the actual analysis of the strength of the correlation was taken from the correlation in the preferred and orthogonal directions as defined below.

To determine the preferred axis of the spatially modulated cells, we calculated the correlation coefficients between firing rate and position along 180 different axes evenly spaced on the half circle and determined the one for which the value of the correlation coefficient was maximal (*Θ*_*max*_, the preferred axis) and the one for which the absolute value of the correlation coefficient was minimal (*Θ*_*min*_). As expected, *Θ*_*min*_ was tightly distributed around the orthogonal axis of *Θ*_*max*_. To simplify the analysis, we used *Θ*_*max*_ ± 90^*o*^ as the null axis for all neurons.

To evaluate the axial tuning similarity of two sequential recording sessions (examples presented in Figure 4 G and O), we calculated the correlation coefficient between the two tuning curves of firing rate to position along *Θ*_*max*_ and compared it to the correlation coefficient of the tuning curve from the first session to the tuning curves of 5000 shuffled spike trains obtained from the second recording session. Comparing these yielded a p-value estimating the probability of obtaining axial-tuning curves with greater similarity than of the cell by chance.

### Axiality tests

To determine whether the spatially modulated cells were tuned to the preferred direction regardless of position along the null direction (i.e., axial encoding rather than Gaussian encoding in the form of place cells centered near the boundaries of the environment), we tested three alternative models for the modulation of firing rates in space: a sigmoid model assuming axial modulation, a Gaussian model assuming local modulation, and a uniform model of no modulation (control).

We tested our ability to differentiate these alternatives on simulated data (Supplementary Figure 3). Each model was tested with three different trajectories from those where spatially modulated cells were recorded in our data. For the sigmoid model, we simulated 500 axially-tuned firing patterns generated from three real tuning curves of firing rate to position as recorded in our data. These were rotated to random preferred directions in space, while in the null (orthogonal) direction the tuning of rate to position was bounded to a uniform distribution. Examples are presented in Supplementary Figure 3 A and D. Each simulated firing pattern was then fitted with a two-dimensional Gaussian (Supplementary Figure 3 B and E, respectively) and with a sigmoid (Supplementary Figure 3 C and F, respectively).

We used the Matlab **fit** function to minimize the least square difference from the sigmoid:

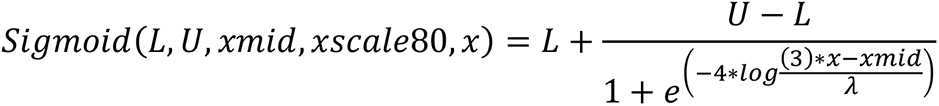

where *L* and *U* are the lower and upper asymptotes of the sigmoid, respectively, *xmid* is the point where the curve goes through 50% of the transition, and*λ* is the width in which 80% of the transition occurs. The parameter *x* represents position along the *Θ*_*max*_ axis of the simulated cell.

The quality of the fit was assessed as the correlation between the rate map of the simulated data and the rate map yielded by the fit model. Each simulated dataset was also fitted with a Gaussian using the Matlab **fitgmdist** function and goodness of fit was assessed in the same way.

Simulated firing rates for the Gaussian model were generated by choosing Gaussians whose centers were within 10 cm of the edges of the environment. This simulated the near border tuning present in our recordings and provided an alternative to the sigmoid tuning. The same pipeline was used for this model as for the former. An example of a simulated Gaussian encoding cell and rate maps fitted from the two models is presented in Supplementary Figure 3 G-I. We also repeated the process using 500 rate maps generated from a uniform distribution (Supplementary Figure 3 J-L).

The results of the simulation (Figure 5B) showed that axial encoding cells could be distinguished from Gaussian encoding cells using the correlation coefficients between the cell’s rate map and the rate map from the fit models. For this purpose, we drew threshold lines which created a criterion for axially tuned cells with very few false positives. That is, every neuron with a correlation to the sigmoid model above 0.7 and above the correlation to the Gaussian model was considered to be axial encoding. Only 4 of the 500 Gaussian-tuned simulated cells and 7 of the non-tuned (uniform) simulated cells were thus classified as axial encoding (a false positive rate of around 0.011). In addition, 36 of the 500 axially-tuned simulated cells were classified as non-axial (a false negative rate of around 0.072). We applied the thresholds derived from the simulations to the corresponding results for the real data to categorize the actual neurons into axial encoding or non-axial encoding (Figure 5A).

## Acknowledgments

We are grateful to Jacob Vecht from Deuteron Technologies Ltd. and Tal Novoplansky-Tzur for helpful technical assistance. We gratefully acknowledge financial support from THE ISRAEL SCIENCE FOUNDATION—FIRST Program (Grant no. 281/15), THE ISRAEL SCIENCE FOUNDATION—FIRST Program (Grant no. 555/19), THE ISRAEL SCIENCE FOUNDATION (Grant no. 211/15), The Human Frontiers Science Foundation Grant RGP0016/2019, and the Helmsley Charitable Trust through the Agricultural, Biological and Cognitive Robotics Initiative of Ben-Gurion University of the Negev.

## Author contributions

Conceptualization, L.C., E.V., O.D., R.S., methodology, L.C., E.V., O.D., R.S., formal analysis, L.C., E.V., O.D., R.S., investigation L.C., E.V., O.D., R.S, writing, L.C., E.V., O.D., R.S.

---

**Supplementary Videos 1-3 | Video Examples of Spatially Modulated Cells.** Left panels-a video recording shows an example of a spatially modulated cell wirelessly recorded from the goldfish brain while freely exploring the experimental water tank. Spiking activity can be heard throughout the audio channel. Right panels-swimming trajectories (black curves, top panels) and spiking locations (red dots) together with the occupancy corrected rate maps (bottom panels) of the cell presented in the corresponding video. Examples correspond to the cells presented in Figure 2 A-B (Supplementary Video 1), Supplementary Figure 1C (Supplementary Video 2) and Supplementary Figure 1A (Supplementary Video 3).

**Supplementary Figure 1.**
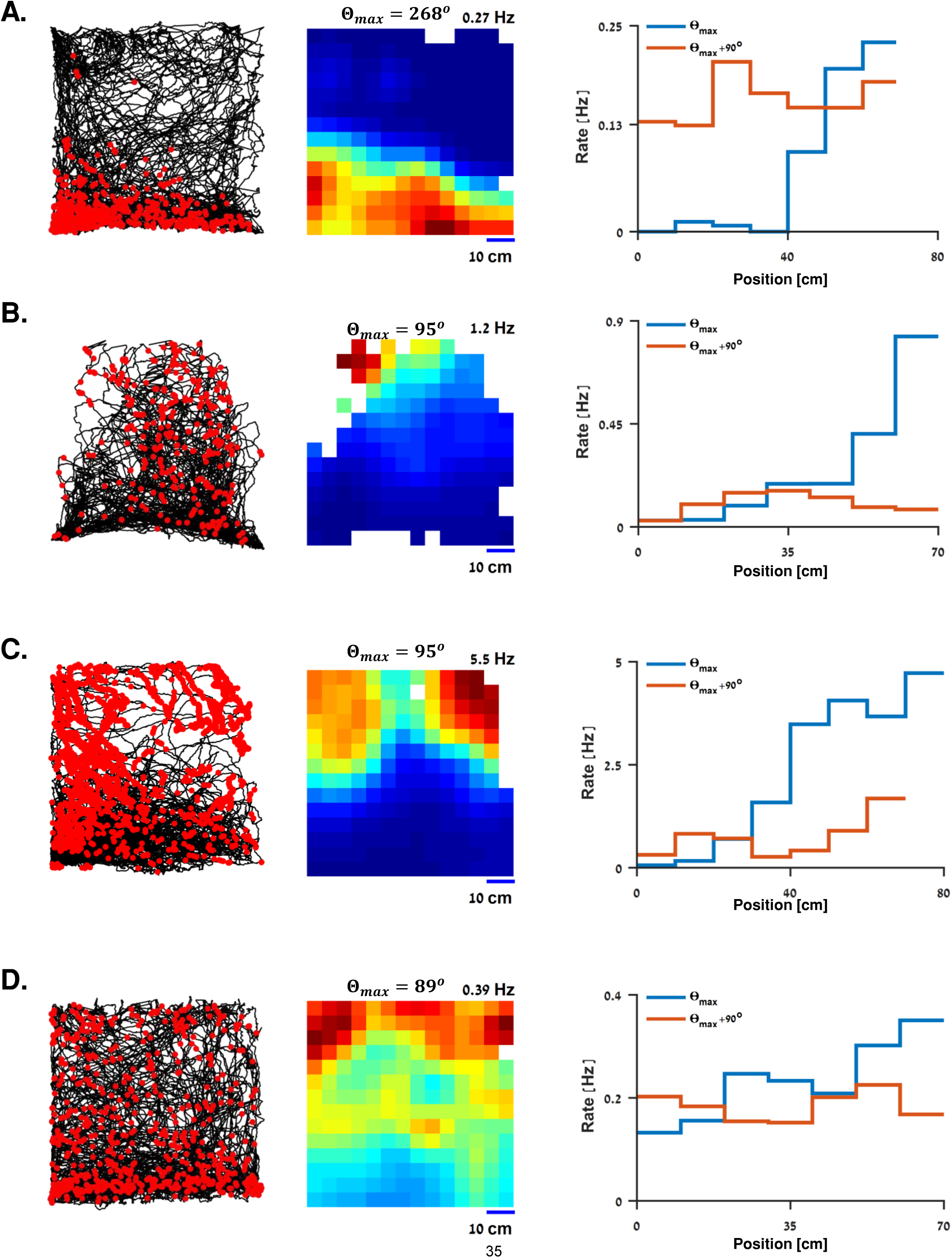
Additional examples spatial modulations along the vertical axis. Each panel in A-D represents a single cell. For each cell the graph shows the spiking activity (red dots, left panels) over the fish’s trajectory (black curves), the color-coded rate map (middle panels) ranging from dark blue (zero) to dark red (maximal firing rate) and the tuning curves (right panels) of the firing rate to position of the fish along the preferred axis (*Θ*_*max*_, blue curve) and its orthogonal axis (*Θ*_*max*_ + 90^*o*^, orange curve). Gradual tuning is shown for the *Θ*_*max*_ axis.

**Supplementary Figure 2.**
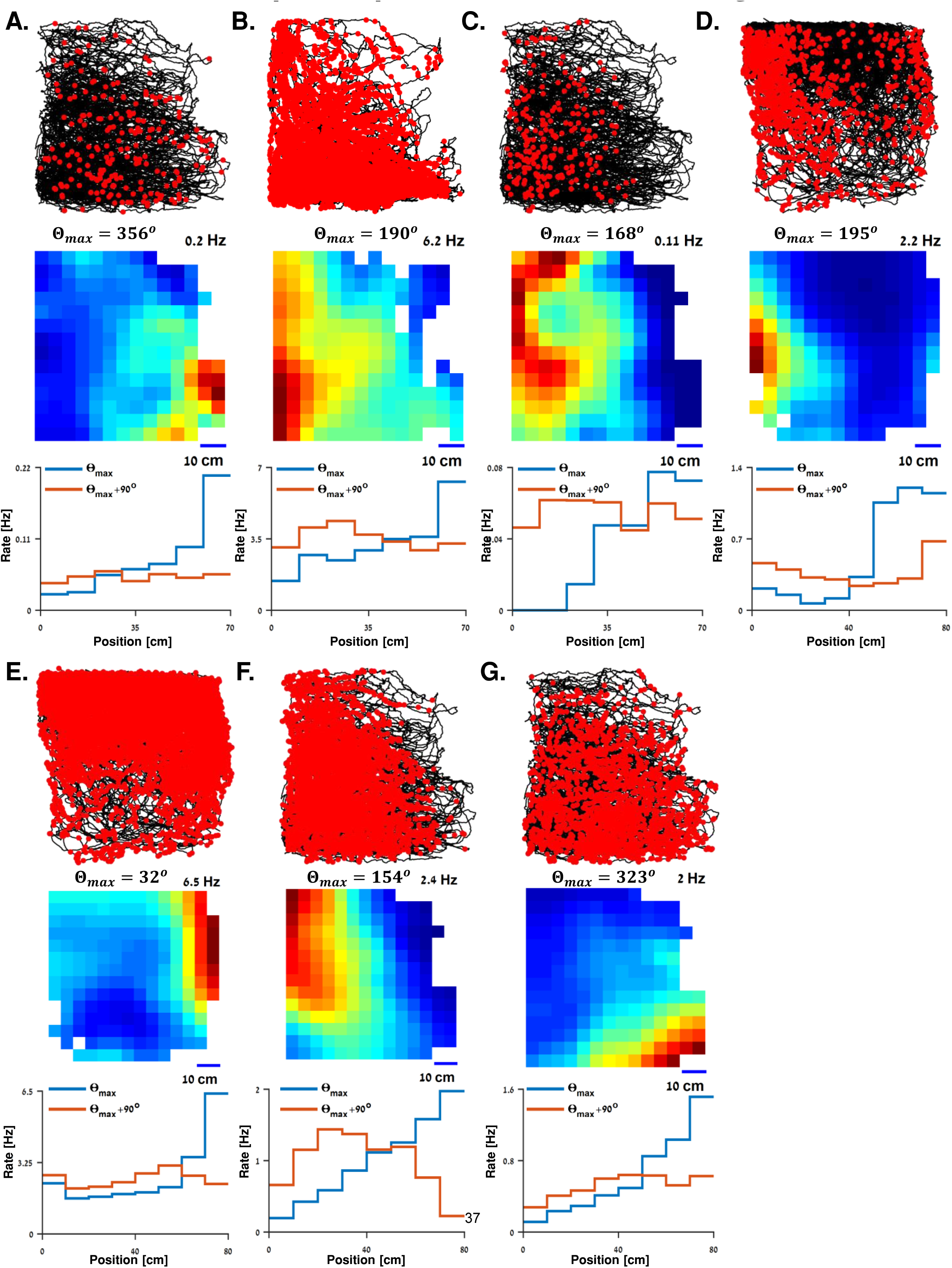
Additional examples-spatial modulations along other axes. **A-D.** Examples of cells with preferred tuning to the horizontal axis. For each cell, the graphs show the spiking activity (red dots, top panels) over the fish’s trajectory (black curves), the color-coded rate map (middle panels) ranging from dark blue (zero) to dark red (maximal firing rate) and the tuning curves (bottom panels) of firing rate to position of the fish along the preferred axis (*Θ*_*max*_, blue curve) and its orthogonal axis (*Θ*_*max*_ + 90^*o*^, orange curve). Gradual tuning is shown for the *Θ*_*max*_axis**. E-G.** Examples of cells with preferred tuning to other axes in space. For each cell the graph shows the spiking activity (red dots, top panels) over the fish’s trajectory (black curves), the color-coded rate map (middle panels) ranging from dark blue (zero) to dark red (maximal firing rate) and the tuning curves (bottom panels) of firing rate to the projection of position of the fish along the preferred axis (*Θ*_*max*_, blue curves) and its orthogonal axis (*Θ*_*max*_ + 90, orange curves). Gradual tuning is shown for the *Θ*_*max*_axis.

**Supplementary Figure 3.**
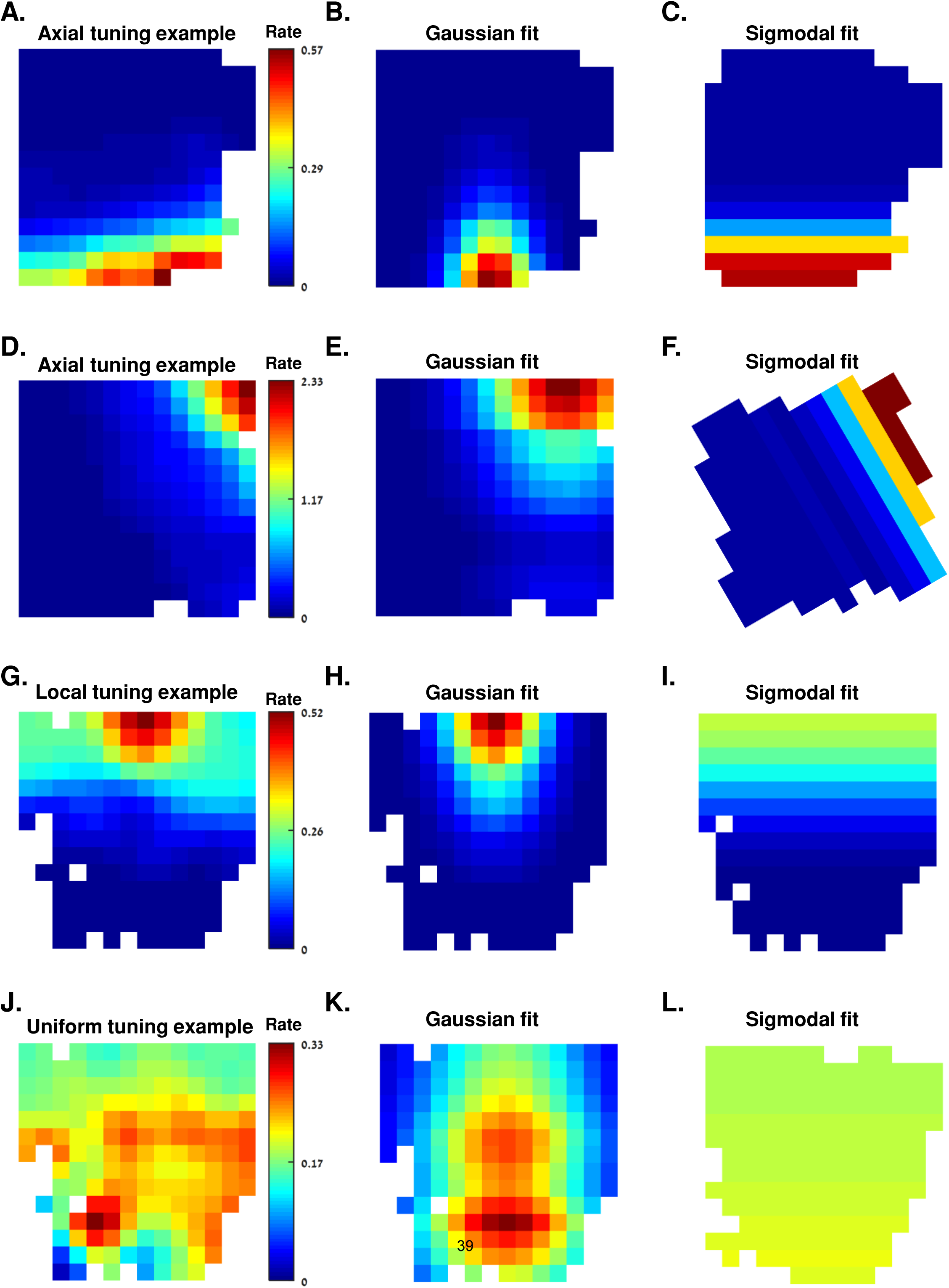
Examples of classification of simulated cells into axial encoding, local encoding, and non-encoding. Three groups of cells (500 cells in each group) were simulated with different tuning properties in space. For each group, two models were tested to characterize the resulting firing pattern. **A.** Example of a simulated rate map with axial tuning along a major axis in space. Same color-bar was used for the rate maps in panels B and C. **B.** A rate map fitted to the rate map in A using 2DGaussian tuning. **C.** A rate map fitted to the rate map in A using sigmoid tuning to the preferred axis of the cell in A. **D-F.** Another example of a simulated rate map with axial tuning along an axis that was not parallel to a Cartesian axis. **G-I.** Another simulated example with a firing pattern locally tuned near a boundary in the environment. **J-L.** Another simulated example with a uniformly distributed firing pattern in space.

**Supplementary Figure 4.**
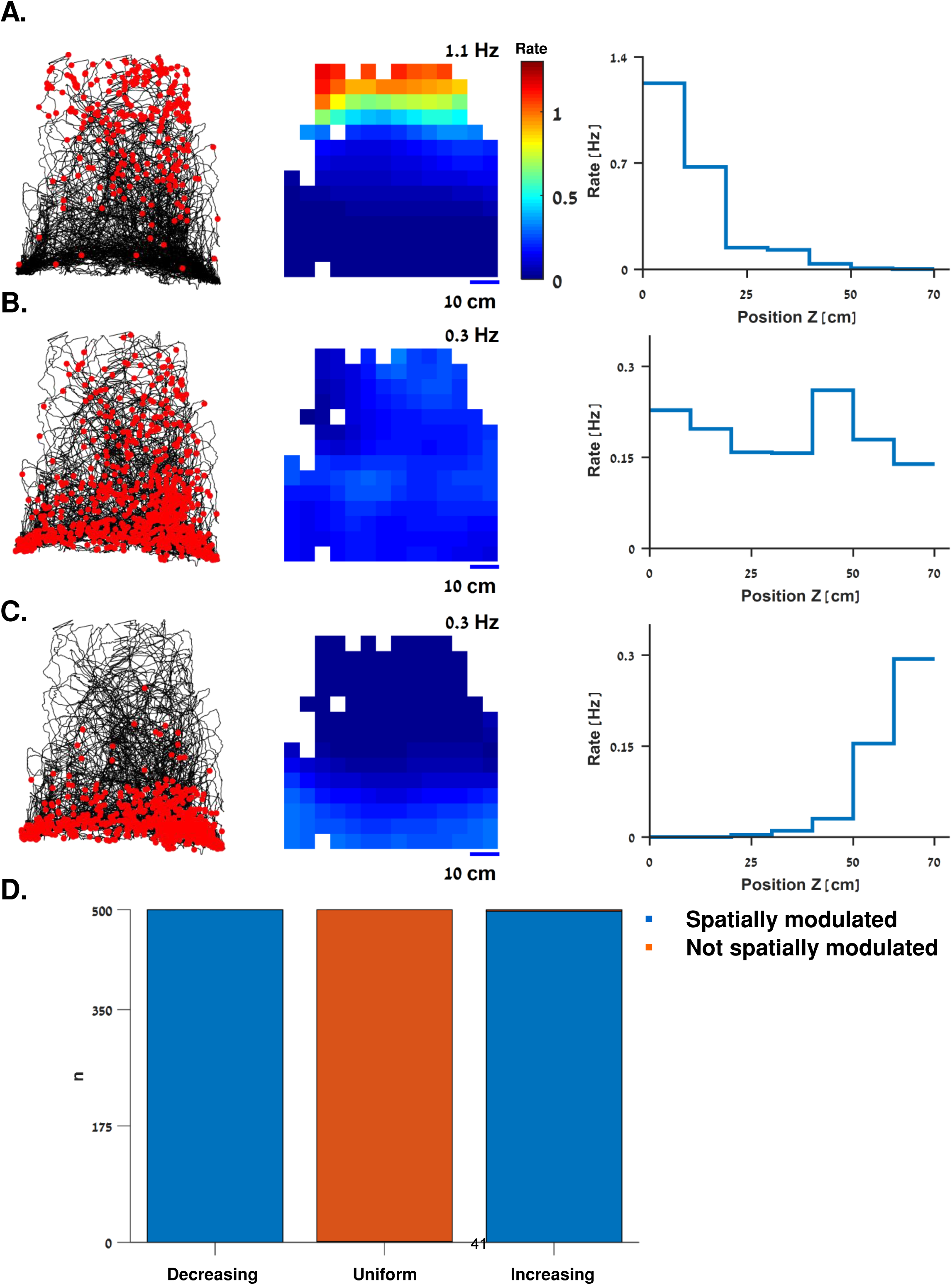
Validating the stability of the analysis against a non-uniform coverage of the fish trajectory. We used three tuning curves of firing rate to position in the vertical axis as recorded from three different fish to simulate firing patterns in a real trajectory (panel A, left panel, black curve) which partially covered the experimental water tank. This trajectory corresponds to the cell presented in Supplementary Figure 1B. For each tuning curve, we simulated 500 spike trains. **A-C**. examples of spiking patterns (red dots, left panels) over the swimming trajectory (black curves) together with a color-coded occupancy-corrected heatmap (middle panels, same color bar as in panel A) and the tuning curve used to simulate them (right panels). The tuning curves were gradually decreased (A), were quasi-uniformly distributed (B) or gradually increased (C) with position along the vertical axis. **D.** Simulation results. Each simulated spike train was then tested to determine whether it crossed the threshold for a spatially modulated cell (see *Materials and Methods*). Out of the 1500 spike trains, 1497 were classified correctly (false negative rate<0.002).

**Supplementary Figure 5.**
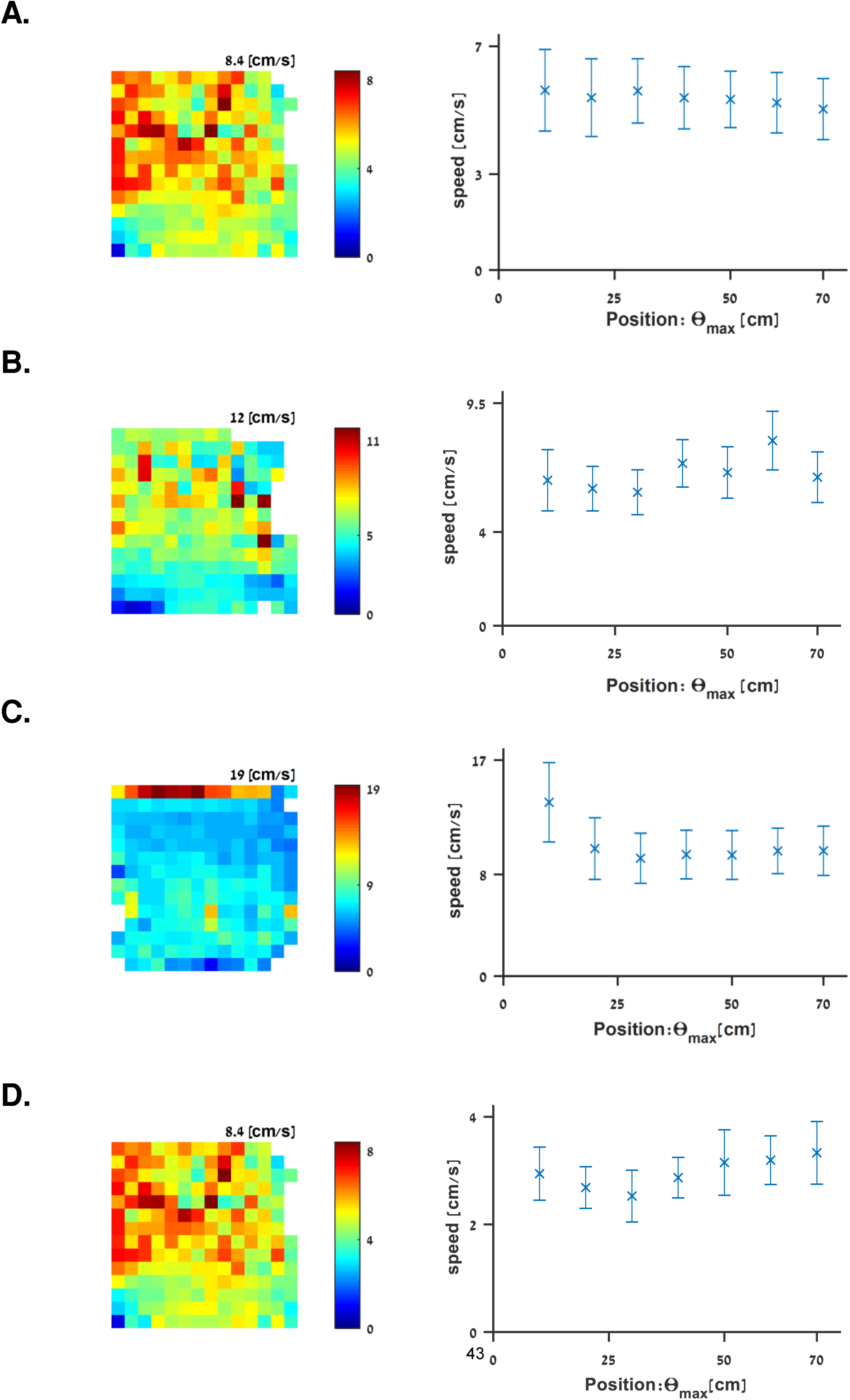
Speed distribution across the water tank. Examples of swimming speed distributions of spatially modulated cells. For each cell, a swimming speed map (left panels) is presented, color coded from dark blue (zero) to dark red (maximal swimming speed, indicated on the top right side of each panel). Also shown are the error bars (mean and standard error) of speed across position along the preferred axis of the corresponding cells (right panels). Aside from the physical limits of swimming near the edges of the tank, no clear monotone pattern was found. Examples correspond to the cells presented in **A.** Figure 2 A-D, **B.** Figure 2 E-H, **C.** Figure 2 I-L and **D.** Figure 2 M-P.

**Supplementary Table 1.**

| **Classified as:** | **Axial** | **Gaussian** | **Uniform** |
| --- | --- | --- | --- |
| **Spatially modulated cells** | **21** | **2** | **4** |
| **Other cells** | **3** | **4** | **36** |

**Supplementary Table 2.**

| **max-p**<br>**Threshold** | **Spatial Information**<br>**Threshold** | **# spatially**<br>**modulated cells** |
| --- | --- | --- |
| **1.00E-03** | **0.05** | **23** |
| **1.00E-03** | **0.1** | **22** |
| **1.00E-03** | **0.2** | **17** |
| **1.00E-03** | **0.3** | **12** |
| **1.00E-02** | **0.05** | **28** |
| **1.00E-02** | **0.1** | **27** |
| **1.00E-02** | **0.2** | **21** |
| **1.00E-02** | **0.3** | **15** |
| **0.025** | **0.05** | **32** |
| **0.025** | **0.1** | **31** |
| **0.025** | **0.2** | **24** |
| **0.025** | **0.3** | **17** |
| **0.05** | **0.05** | **36** |
| **0.05** | **0.1** | **35** |
| **0.05** | **0.2** | **27** |
| **0.05** | **0.3** | **20** |

## References

1. Hafting T, Fyhn M, Molden S, Moser M-B, Moser EI. Microstructure of a spatial map in the entorhinal cortex. Nature Publishing Group; 2005;436(7052): 801–806.

2. Baxendale S, Whitfield TT. Zebrafish inner ear development and function. Elsevier; 2014. 43 p.

3. O’Keefe J, Dostrovsky J. The hippocampus as a spatial map: preliminary evidence from unit activity in the freely-moving rat. Elsevier Science; 1971;

4. John O. Hippocampal Neurophysiology in the Behaving Animal. Oxford University Press; 2006; 475– 548.

5. Cohen L, Vinepinsky E, Segev R. Wireless electrophysiological recording of neurons by movable tetrodes in freely swimming fish. 2019;(153): e60524.

6. Northcutt RG. Connections of the lateral and medial divisions of the goldfish telencephalic pallium. Wiley Online Library; 2006;494(6): 903–943.

7. Agarwal A, Sarel A, Derdikman D, Ulanovsky N, Gutfreund Y. Spatial coding in the hippocampus of flying owls. Cold Spring Harbor Laboratory; 2021;

8. Lipp H-P, Vyssotski AL, Wolfer DP, Renaudineau S, Savini M, Tröster G, et al. Pigeon homing along highways and exits. Elsevier; 2004;14(14): 1239–1249.

9. Rowland DC, Roudi Y, Moser M-B, Moser EI. Ten years of grid cells. Annual Reviews; 2016;39: 19–40. 10.

10. Ben-Yisahay E, Krivoruchko K, Ron S, Ulanovsky N, Derdikman D, Gutfreund Y. Directional tuning in the hippocampal formation of birds. Elsevier; 2021;31(12): 2592–2602. e4.

11. Northcutt RG. Forebrain evolution in bony fishes. Elsevier; 2008;75(2–4): 191–205.

12. Buzsáki G, Moser EI. Memory, navigation and theta rhythm in the hippocampal-entorhinal system. Nature Publishing Group; 2013;16(2): 130–138.

13. Lever C, Burton S, Jeewajee A, O’Keefe J, Burgess N. Boundary vector cells in the subiculum of the hippocampal formation. Soc Neuroscience; 2009;29(31): 9771–9777.

14. Grieves RM, Jedidi-Ayoub S, Mishchanchuk K, Liu A, Renaudineau S, Jeffery KJ. The place-cell representation of volumetric space in rats. Nature Publishing Group; 2020;11(1): 1–13.

15. Payne HL, Lynch GF, Aronov D. Neural representations of space in the hippocampus of a food-caching bird. American Association for the Advancement of Science; 2021;373(6552): 343–348.

16. Wehner R. HOW MINIATURE BRAINS SOLVE COMPLEX TASKS. World Scientific; 2001. 8 p.

17. Peter RE, Gill VE. A stereotaxic atlas and technique for forebrain nuclei of the goldfish, Carassius auratus. Wiley Online Library; 1975;159(1): 69–101.

18. Dittman A, Quinn T. Homing in Pacific salmon: mechanisms and ecological basis. 1996;199(1): 83–91.

19. Taylor GK, Holbrook RI, de Perera TB. Fractional rate of change of swim-bladder volume is reliably related to absolute depth during vertical displacements in teleost fish. The Royal Society; 2010;7(50): 1379–1382.

20. Kato T, Yamada Y, Yamamoto N. Ascending gustatory pathways to the telencephalon in goldfish. Wiley Online Library; 2012;520(11): 2475–2499.

21. Vinepinsky E, Cohen L, Perchik S, Ben-Shahar O, Donchin O, Segev R. Representation of edges, head direction, and swimming kinematics in the brain of freely-navigating fish. Nature Publishing Group; 2020;10(1): 1–16.

22. Broglio C, Gómez A, Durán E, Salas C, Rodríguez F. Brain and cognition in teleost fish. Wiley; 2011. 34 p.

23. Moser EI, Kropff E, Moser M-B. Place cells, grid cells, and the brain’s spatial representation system. Annual Reviews; 2008;31: 69–89.

24. Bass AH, McKibben JR. Neural mechanisms and behaviors for acoustic communication in teleost fish. Elsevier; 2003;69(1): 1–26.

25. Northcutt RG. Do teleost fishes possess a homolog of mammalian isocortex? S. Karger AG; 2011;78(2): 136.

26. Mueller T, Dong Z, Berberoglu MA, Guo S. The dorsal pallium in zebrafish, Danio rerio (Cyprinidae, Teleostei). Elsevier; 2011;1381: 95–105.

27. Vinepinsky E, Donchin O, Segev R. Wireless electrophysiology of the brain of freely swimming goldfish. Elsevier; 2017;278: 76–86.

28. Moser EI, Moser M-B, McNaughton BL. Spatial representation in the hippocampal formation: a history. Nature Publishing Group; 2017;20(11): 1448–1464.

29. Moser EI, Roudi Y, Witter MP, Kentros C, Bonhoeffer T, Moser M-B. Grid cells and cortical representation. Nature Publishing Group; 2014;15(7): 466–481.

30. Fraser PJ, Shelmerdine RL. Dogfish hair cells sense hydrostatic pressure. Nature Publishing Group; 2002;415(6871): 495–496.

31. Ulanovsky N. Neuroscience: how is three-dimensional space encoded in the brain? Elsevier; 2011;21(21): R886–R888.

32. Segev R, Goodhouse J, Puchalla J, Berry MJ. Recording spikes from a large fraction of the ganglion cells in a retinal patch. Nature Publishing Group; 2004;7(10): 1155–1162.

33. Schulz-Mirbach T, Heß M, Metscher BD, Ladich F. A unique swim bladder-inner ear connection in a teleost fish revealed by a combined high-resolution microtomographic and three-dimensional histological study. BioMed Central; 2013;11(1): 1–13.

34. Lewicki MS. A review of methods for spike sorting: the detection and classification of neural action potentials. IOP Publishing; 1998;9(4): R53.

35. Ginosar G, Aljadeff J, Burak Y, Sompolinsky H, Las L, Ulanovsky N. Locally ordered representation of 3D space in the entorhinal cortex. Nature Publishing Group; 2021;596(7872): 404–409.

36. Alme CB, Miao C, Jezek K, Treves A, Moser EI, Moser M-B. Place cells in the hippocampus: eleven maps for eleven rooms. National Acad Sciences; 2014;111(52): 18428–18435.

37. McNaughton BL, Battaglia FP, Jensen O, Moser EI, Moser M-B. Path integration and the neural basis of the’cognitive map’. Nature Publishing Group; 2006;7(8): 663–678.

38. Diogo R. Origin, Evolution and Homologies of the Weberian Apparatus: A New Insight. 2009;27(2). 39.

39. Davis VA, Holbrook RI, de Perera TB. Fish can use hydrostatic pressure to determine their absolute depth. Nature Publishing Group; 2021;4(1): 1–5.

40. Mathis A, Mamidanna P, Cury KM, Abe T, Murthy VN, Mathis MW, et al. DeepLabCut: markerless pose estimation of user-defined body parts with deep learning. Nature Publishing Group; 2018;21(9): 1281–1289.

41. Solstad T, Boccara CN, Kropff E, Moser M-B, Moser EI. Representation of geometric borders in the entorhinal cortex. American Association for the Advancement of Science; 2008;322(5909): 1865–1868.

42. Altman JA, Butusov MM, Vaitulevich SF, Sokolov AV. Responses of the swimbladder of the carp to sound stimulation. Elsevier; 1984;14(2): 145–153.

43. Tsoar A, Nathan R, Bartan Y, Vyssotski A, Dell’Omo G, Ulanovsky N. Large-scale navigational map in a mammal. National Acad Sciences; 2011;108(37): E718–E724.

44. Holbrook RI, de Perera TB. Separate encoding of vertical and horizontal components of space during orientation in fish. Elsevier; 2009;78(2): 241–245.

45. de Perera TB, Holbrook R, Davis V, Kacelnik A, Guilford T. Navigating in a volumetric world: metric encoding in the vertical axis of space. Cambridge University Press; 2013;36(5): 546.

